# Subcellular, biochemical and biophysical alterations in two glial cell models of ARSACS

**DOI:** 10.1101/2024.04.15.589510

**Authors:** Fernanda Murtinheira, Ana Sofia Boasinha, João Belo, Luana Macedo, Elisa Farsetti, Tiago T. Robalo, Vukosava M. Torres, Francisco R. Pinto, Adelaide Fernandes, Patricia Nascimento, Mario S. Rodrigues, Federico Herrera

## Abstract

Autosomal recessive spastic ataxia of Charlevoix-Saguenay (ARSACS) is a developmental and degenerative disorder caused by loss-of-function mutations in the gene that codifies for the sacsin chaperone. Sacsin was initially described as a neuronal protein but is found in various cell types, including astroglial, microglial, kidney, and skin cell lines. We and others have shown that virtually all cell and animal models of ARSACS show disruption of intermediate filament (IF) cytoskeleton and organelle distribution. This article extends previous observations from our lab on the C6 astroglial-like model and describes the development and characterization of a new human microglial cell model based on HMC3 cells. HMC3 cells knocked out for sacsin show similar alterations to C6 astroglial-like cells: aberrant distribution of IF network and organelles, downregulation of developmental transcription factors STAT3 and Smad1, and alterations in autophagy machinery and markers of intracellular stress, resulting in morphological changes. Our results extend previous observations suggesting a possible role for glial cells in ARSACS and provide new tools to understand the glial-specific mechanisms involved in this pathology.

## Introduction

Loss-of-function mutations in the sacsin chaperone cause autosomal recessive spastic ataxia of Charlevoix-Saguenay (ARSACS), a developmental and degenerative disorder characterized by diverse motor symptoms (Larivière et al., 2015; Vermeer et al., 2020). Although it is generally described as a neuronal protein, high levels of sacsin expression have been found in many other cell types, such as astroglial, microglial, kidney and skin cell lines. The Protein Atlas, Harmonizome and BrainRNASeq public databases indicate that medium- high sacsin mRNA levels can be found in mouse and human astrocytes, Müller glia, microglia, and macrophages. Mouse sacsin mRNA expression can be as high in astrocytes as in neurons, displaying the highest levels in younger animals (postnatal day 7) and decreasing with age (Clarke et al., 2018). Human fetal astrocytes express the same mRNA expression levels as neurons, also decreasing as astrocytes mature (Zhang et al., 2016). We have recently confirmed experimentally that primary rat astrocytes express high levels of the sacsin protein, and so did C6 (glioblastoma, astroglial-like) and N9 (microglial) rodent cell lines (Murtinheira et al., 2022). There are several indicators of neuroinflammation in ARSACS (Bondio et al., 2023), suggesting a possible role for glial cell dysfunction in ARSACS. Furthermore, ARSACS patients show signs of premature ageing (i.e. lipofuscin aggregates) in skin biopsies (Stevens et al., 2013), and ARSACS-derived fibroblasts show important cellular and molecular alteration (Bradshaw et al., 2016), suggesting that sacsin biological roles go beyond the central nervous system.

Disruption of intermediate filament (IF) cytoskeleton and of organelle distribution are largely the most common hallmarks in all cellular and animal models of ARSACS (Dabbaghizadeh et al., 2022; Gentil et al., 2018; Murtinheira et al., 2022), including nerve and glial cells. IFs play an important role in maintaining the mechanical and viscoelastic properties of cells (Sivaramakrishnan et al., 2008), which are important for a wide range of cellular functions, such as cell migration, division, and differentiation. IFs provide structural support and help to maintain cell and tissue shape, especially under mechanical stress or strain (Pogoda & Janmey, 2023). Additionally, intermediate filaments also act as scaffolds involved in cell signalling and in the localization and traffic of organelles, such as mitochondria, lysosomes and other vesicles. Alterations in IF networks are associated with at least 80 different human pathologies, some of them with impact on neural functions similar to ARSACS (Omary, 2009), such as Alexandeŕs disease (AxD) and Giant Axonal Neuropathy (GAN).

In this article, we extend our previous observations on the C6 astroglial-like model of ARSACS (Murtinheira et al., 2022), reporting a series of biochemical and biophysical alterations that could be relevant to glial dysfunction in ARSACS. Additionally, we develop a new human microglial cell model of ARSACS and confirm part of the alterations observed in C6 cells. Our results further support a possible role for glial cells in ARSACS.

## Methods

### 1. Cell Cultures

C6 (rat, glioblastoma, astroglial-like) and HMC3 (human, microglial) cells were acquired from ATCC (references CCL-107 and CRL-3304, respectively). Cells were maintained in DMEM medium supplemented with 10% v/v Fetal Bovine Serum (FBS), 1% L-glutamine and 1% Penicillin and Streptomycin mix and maintained at 37°C in a 5% CO2 atmosphere. Sacsin knockout was achieved as previously described, using a CRISPR/Cas9 tool commercially available (Santa Cruz Biotechnologies) and a FACSAria III cell sorter (BD Biosciences) (Murtinheira et al., 2022). Cells were plated on different sterile plastic dishes or on sterile glass coverslips and allowed to adhere for 16–24 h before experiments and/or sample preparation. When indicated, cells were transfected 24h after seeding with mammalian expression plasmids to visualize Golgi (EYFP-Golgi7, Addgene plasmids # 56590), using JetPRIME transfection reagent (Polyplus transfection, Illkirch, France) at 1:3 transfection ratios. HMC3 and HMC3^Sacs-/-^ cells were incubated with 1mg/ml of myelin debris previously stained with BASHY probe as described (Pinto et al., 2021) for 1h to mimic microglia neuroprotective ability to remove debris, or with 0.0025% (v/v) 1 mm zymogen-coated fluorescent latex beads (Sigma Chemical) for 75 min to mimic microglia reactivity against infections. Mitochondria were visualized by incubating cells with Mitoview^TM^ Fix 640 (Biotium; Fremont, CA, USA) for 2h before imaging.

#### 1.1. Cell viability assays

For MTT assays, C6 and C6^Sacs−/−^ cells were seeded onto 96-well plates at a concentration of 10^4^ cells/well in a standard growth medium for 24 hours. After washing with PBS, they were incubated in serum-free medium for 48 hours.Then, cells were incubated with MTT (0.5 mg/mL) for 2 h. The medium was then replaced with DMSO (100% *v*/*v*). After 15 min of incubation in the dark at RT, absorbance was determined using an automatic microplate reader (Tecan Sunrise Microplate Reader) (Tecan, Männedorf, Switzerland) at 595 nm.

#### 1.2. Intracellular ROS assays

Cells were seeded in 6-well plates (5 × 10^5^ cells/well). After 48 hours in serum-free medium, cells were detached by trypsin (0.05% *w/v*), collected, resuspended in PBS, and incubated with 10 μM DCFH-DA (All reactive oxygen species) in DMEM medium (without serum) for 20 min at 37 °C in the dark. After washing cells twice with PBS, pellets were resuspended in PBS with DAPI (1 µg/mL) to discriminate between live and dead cells, and fluorescence was immediately analyzed by means of a BD LSRFortessa X-20 cell analyzer (BD Biosciences, San Jose, CA, USA). At least 10,000 events (low velocity) were recorded for analysis with FLOWJO software Version 9 (Emerald Biotech Co., Ltd., Córdoba, Spain).

#### 1.3. Cell cycle

C6 and C6^Sacs-/-^ cells were seeded in 6-well plates (5 × 10^5^ cells/well) in a standard growth medium for 24 hours. Afterwards, they were washed with PBS and exposed to a serum-free medium for 48 hours. For cell cycle analysis, floating and adherent cells were collected by centrifugation and fixed by dropwise adding ice-cold 70% ethanol and incubating overnight at 4°C. Cells were centrifuged at 500 x g for 5 min, washed twice with PBS, and resuspended in a propidium iodide solution (50 μg/ml in PBS) containing RNase A (100μg/ml). After 20 min of incubation at 37°C, cells were analyzed using a FACSCalibur flow cytometer (BD Biosciences, Franklin Lakes, NJ, USA). Ten thousand events were recorded for cell cycle analysis with FLOWJO software Version 9 (Emerald Biotech Co., Ltd., Córdoba, Spain).

### 2. Microscopy

Cells grown on coverslips on 24-well plates were either live-imaged or fixed for immunocytochemistry. Cells were washed with Dulbecco’s Phosphate Buffered Saline (DPBS; San Marcos, TX, USA) and fixed with methanol for 15 min at -20°C. Cells were then washed with DPBS (3 x 5 minutes) and permeabilized with 0,1% Triton X-100 in DPBS for 10 minutes. After 3 washes with DPBS, cells were blocked with 1% BSA in DPBS-T for 1h at room temperature. Afterwards, incubation with primary antibodies (Table 1) was carried out overnight at 4°C in a dark wet chamber. Cells were washed with DPBS for 5 minutes and incubated with the corresponding fluorochrome-conjugated secondary antibodies (Table 2) for 2h at room temperature. Nuclei were counterstained with Hoechst 33342 (Molecular Probes, Willow Creek Rd Eugene, OR, USA). The coverslips were mounted on microscopy slides and imaged using a Leica DMI4000B widefield Microscope, equipped with a x63 oil objective HCX PL APO and a Leica DFC365 FX 1.4MP CCD camera (6.45×6.45 μm/pixel). Images were analyzed with the ImageJ/Fiji software (Rasband, W.S., ImageJ, U. S. National Institutes of Health, Bethesda, Maryland, USA, https://imagej.net/ij/, 1997-2018.). To assess microglia morphology, following fixation cells were incubated with phalloidin (Molecular Probes, dilution 1:400) for 30 minutes, to stain actin filaments in the cells, and nuclei counterstained with DAPI (SIGMA, dilution 1:1000 in PBS) for approximately 5 minutes. Fluorescence images were obtained by confocal microscopy using Leica DMi8-CS inverted microscope with Leica LAS X software and different z-stacks were analyzed with AIVIA software (Microscopy image analysis software) to determine cell morphology parameters and phagocytosis.

**Table 1:** Primary antibodies.

| Primary antibodies |  |  |  |  |
| --- | --- | --- | --- | --- |
| Antibody | Dilution | Manufacture | City, Country | Reference |
| Rabbit polyclonal anti- Sacsin | 1:1000 | MilliPore | Burlington, MA, USA | ABN1019 |
| Mouse monoclonal anti-Sacsin (G-3) | 1:1000 | Santa Cruz Biotechnology | Dallas, TX, USA | Sc-515118 |
| Rabbit polyclonal anti- Glial Fibrillary Acid Protein (GFAP) | 1:1000 | MilliPore | Burlington, MA, USA | AB5804 |
| Mouse monoclonal anti-Plectin (10F6) | 1:1000 | Santa Cruz Biotechnology | Dallas, TX, USA | Sc-33649 |
| Mouse monoclonal anti-Vimentin (3CB2) | 1:1000 | Santa Cruz Biotechnology | Dallas, TX, USA | Sc-80975 |
| Mouse monoclonal anti-Nestin (Rat-401) | 1:1000 | Santa Cruz Biotechnology | Dallas, TX, USA | Sc-33677 |
| Mouse monoclonal anti-STAT3 (F-2) | 1:1000 | Santa Cruz Biotechnology | Dallas, TX, USA | Sc-8019 |
| Rabbit monoclonal anti-STAT3 (D3Z2G) | 1:1000 | Cell Signaling Technology | Danvers, MA, USA | #12640 |
| Mouse monoclonal anti-GAPDH (6C5) | 1:2000 | Santa Cruz Biotechnology | Dallas, TX, USA | Sc-32233 |
| Rabbit polyclonal anti- SMAD1 | 1:1000 | Cell Signaling Technology | Danvers, MA, USA | #9743 |
| Mouse monoclonal anti-GLRX3 (Picot) | 1:1000 | Santa Cruz Biotechnology | Dallas, TX, USA | Sc-390068 |
| Mouse monoclonal anti-E6-AP (Ube3A) | 1:1000 | Santa Cruz Biotechnology | Dallas, TX, USA | Sc-166688 |
| Mouse monoclonal anti-NQO1 | 1:1000 | Santa Cruz Biotechnology | Dallas, TX, USA | Sc-32793 |
| Mouse monoclonal anti- MAP LC3$\alpha/\beta$ | 1:1000 | Santa Cruz Biotechnology | Dallas, TX, USA | Sc-16756 |

**Table 2:**
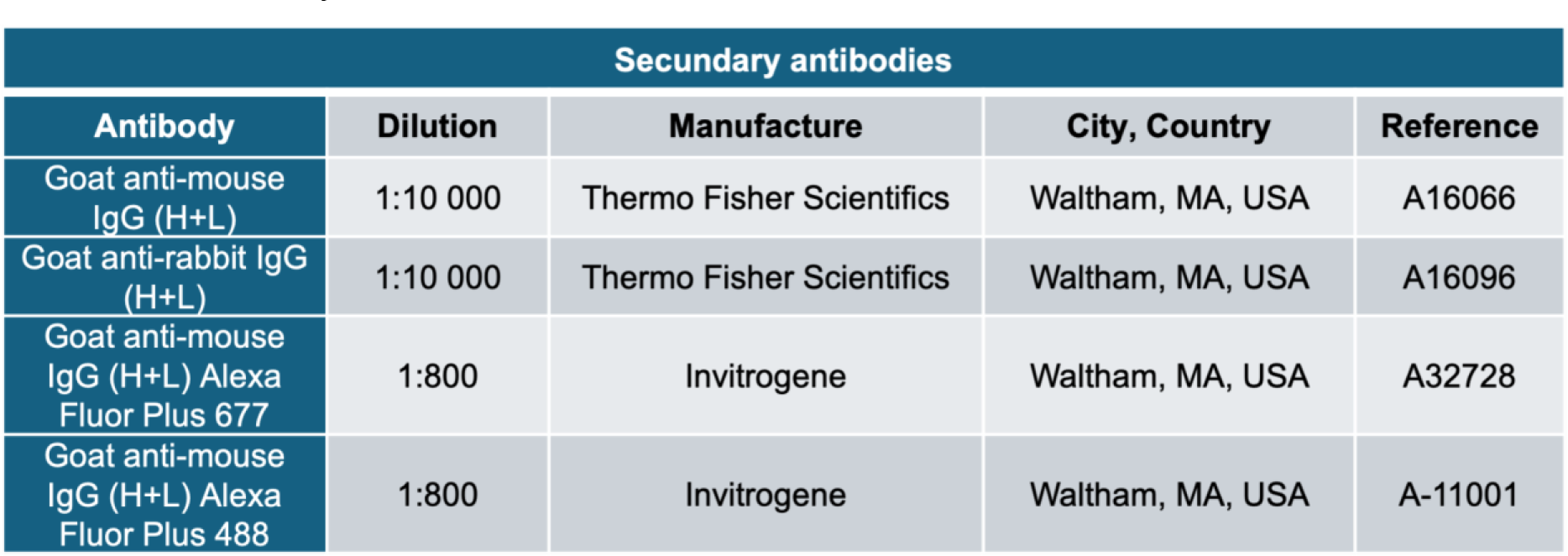
Secondary antibodies.

| Secondary antibodies |  |  |  |  |
| --- | --- | --- | --- | --- |
| Antibody | Dilution | Manufacture | City, Country | Reference |
| Goat anti-mouse IgG (H+L) | 1:10 000 | Thermo Fisher Scientifics | Waltham, MA, USA | A16066 |
| Goat anti-rabbit IgG (H+L) | 1:10 000 | Thermo Fisher Scientifics | Waltham, MA, USA | A16096 |
| Goat anti-mouse IgG (H+L) Alexa Fluor Plus 677 | 1:800 | Invitrogene | Waltham, MA, USA | A32728 |
| Goat anti-mouse IgG (H+L) Alexa Fluor Plus 488 | 1:800 | Invitrogene | Waltham, MA, USA | A-11001 |

### 3. Total protein extraction

Cells seeded in 35mm dishes were washed with DPBS (Cytiva; Marlborough, MA, USA) after the different experimental conditions, and lysed using NP-40 lysis buffer (150 mM NaCl, 50 mM Tris-HCl pH 7.4/7,5, 1% NP-40) or Native lysis buffer (150 mM NaCl, 50 mM Tris-HCl pH 7.4/7.5), both supplemented with cocktails of protease inhibitors (Protease Inhibitor cocktail EDTA free, Abcam, Cambridge, UK) and phosphatase inhibitors (Halt Phosphatase Inhibitor Single-use cocktail, Thermo Fisher Scientifics, Waltham, MA, USA). Cells were scraped from the surfaces of the plates into microcentrifuge tubes and incubated 10 min in ice. A UP200s sonicator (Hielscher Ultrasonics GmbH, Teltow, Germany) was used to disrupt cell membranes for 10 s to release intracellular proteins. Cells were then centrifuged at 10,000 xg for 10 min at 4 °C and the soluble protein fraction was collected. Protein concentration was quantified by the Bradford method on a 96-well plate (Orange Scientific; Braine-l’Alleud, Belgium). A standard curve with known concentrations of bovine serum albumin (BSA, 0.125 to 2 μg/μL) was used to determine protein concentration. Samples were incubated with 200 μL of Bradford solution (Alfa Aesar, Ward Hill, MA, USA) for 5 min and then read at 595 nm in the absorbance microplate reader Sunrise (Tecan, Männedorf, Switzerland).

### 4. Western blotting

Protein extracts (30 µg) were mixed with 4x denaturing loading buffer (0.125 mM Tris pH 6.8; 4% sodium dodecyl sulphate (SDS), 20% glycerol, 10% β-mercaptoethanol, 0.004% bromophenol blue) and boiled for 5 min at 95 °C, followed by incubation in ice for 5 min. Samples were separated by SDS-PAGE in 10% w/v acrylamide gels or in a gradient of 6% + 15% w/v acrylamide gels in running buffer (25 mM Tris-Base; 0,1% SDS; 190 mM Glycine) with a constant voltage of 120 V for 80 min. Proteins were transferred to nitrocellulose membranes (Cytiva, Marlborough, MA, USA) at 100 V for 1 h in transfer buffer (25 mM Tris-Base; 190 mM Glycine; 20% methanol). Membranes were stained with 0.1% w/v Ponceau S (Amresco, Solon, OH, USA) and 5% (v/v) acetic acid to confirm transfer efficiency. Ponceau S solution was washed with Tris-buffered saline (TBS, 150 mM NaCl; 50 mM Tris-base pH 7.4) with 0,05% Tween 20 (TBS-T). Then, membranes were blocked with 5% (w/v) non-fat dry milk in 1x TBS for 1 hour at room temperature. Membranes were washed with TBS-T (3x 10 min.) and incubated with the corresponding primary antibody (Table 1) overnight at 4 °C. Next day, membranes were washed with TBS-T (3x 10 min.) and incubated with the corresponding secondary antibody (1:10000, Table 2) in blocking solution for 2 hours and washed with TBS- T (3x 10 min.). Chemiluminescence detection was performed using the Pierce ECL Plus Western Blotting Substrate and the Amersham Imager 680 blot and gel imager (Cytiva, Marlborough, MA, USA). The integrated intensity of each band was calculated using computer-assisted densitometry analysis with ImageJ software (Rasband, W.S., ImageJ, U. S. National Institutes of Health, Bethesda, Maryland, USA, https://imagej.net/ij/, 1997-2018.).

### 5. Proteomics

#### 5.1. Filter aided sample preparation (FASP digestion)

FASP digestion was carried out as described elsewhere (Wiśniewski et al., 2009), with some minor modifications. Briefly, aliquot of cellular extract, with initial protein input of 20 μg, was diluted to 200 μL with reconstitution buffer (8M Urea, 100mM ammonium bicarbonate - ABC, pH 7.8) and transferred to pre wetted Nanosep™ centrifugal filters, with membrane cut- off of 10kDa (Omega™ - modified polyethersulfone). Buffer exchange was performed adding 200 μL of fresh reconstitution buffer, in five centrifugal cycles (12000 rpm, 5 min.), discarding the flow through fraction. Subsequently, 6 μL of 100mM DTT/100mM ABC was added to the samples and incubated for 50 min. at 56°C, 600 rpm. Reduction was followed by alkylation adding 20 μL of 500 mM IAA/100mM ABC (30 min. at dark). The reconstitution buffer was replaced with 100 mM ABC in four centrifugal cycles (12000 rpm, 5 min.) maintaining the buffer volume at 100 μL. Proteolysis was carried out in-filter, in a wet-chamber at 37°C, during 16h, at 600 rpm, with a mixture of trypsin and LysC (Promega) at an enzyme to substrate ratio of 1:50. Peptides were sequentially eluted by 100 mM ABC (12000 rpm, 5 min.) adding in the last step 0.5% formic acid (FA). Samples were concentrated by means of a vacuum concentrator (SpeedVac, Savant) followed by clean-up with in-house prepared stage tips (AttractSPE® Disks C18, 200 mL tip). Peptides were sequentially eluted (2 x 10 μL) with a solvent mixture of 5% acetonitrile/0.1%FA/ millyQ water, ready for LC/MS analysis.

#### 5.2. nLC-MS/MS analysis

Chromatographic separation was performed on nano Elute™ nano flow ultra high-pressure LC (Bruker, Daltonics) coupled to a ultra-high resolution quadrupole-time-of-flight mass spectrometer (UHR QqTOF, Impact II™ Bruker, Daltonics) with a CaptiveSpray nano booster™ ion source (Bruker, Daltonics). Two microliter of sample was loaded to the Acclaim™ PepMap™ 100 C18 trap column (Thermo Scientific™). Separation was performed on Bruker fifteen™ column (C18, 15 cm × 75 um, 1.9 um, 120 Å) in a 120 min. linear gradient using 0.1% formic acid in MillyQ water (solvent A) and 0.1% formic acid in acetonitrile (solvent B) with constant flow of 300 nl/min in the following regimen: 0 to 115 min., 5 - 35% B; 115 to 118 min., 35 – 95% B; 118-125 min. 95% B. Column was kept at 40°C. The MS acquisition was performed with the following parameters: capillary voltage was set at 1500 V, the drying gas flow rate was 3.0 L/min at a temp of 150 °C, nanobooster at 0.2 bar. The acquisition was performed in the positive ionization mode at spectra rate of 1 Hz, in the range of 150 to 2200 m/z. LC-MS/MS data were acquired using data-dependent (DDA) auto MS/MS method, cycle time 3.0 s, active exclusion was triggered after one spectrum, release after 0.5min.. Reconsider precursor if current intensity/previous intensity is set up to three, and intensity threshold for fragmentation was 2500 counts. The collision energy was tuned between 23–65 eV as a function of the m/z value. The spectra were internally calibrated by post-run infusion of the 1µM sodium formate. Calibration of MS files was performed using Compass Data Analysis 4.4 (Bruker, Daltonics) in high precision calibration mode (HPC) giving error of <1 ppm and standard deviation < 0.2ppm for the mass range between 90 to 1500 m/z.

#### 5.3. Protein identification and label free quantification (LFQ)

The raw files from the mass spectrometer were analyzed using MaxQuant software (ver. No. 2.2.0.0.), as previously reported, using default parameters (Tyanova et al., 2016) [2]. Andromeda search engine was employed for peptide search against the UniProt-SwissProt Rattus Norvegicus database as a reference and contaminants database for common contaminants. Briefly, Trypsin was used as protease with maximum two missed cleavages allowed. The maximum false discovery rate for peptide and protein was 0.01. Main search (parent ion) and peptide (daughter) search tolerances were 20 and 4.5 ppm respectively. The oxidation of methionine and acetylation were used for variable modification while for the fixed modification carbamidomethylation of cysteine was used. The LFQ Label-free quantification (LFQ) was performed using classic normalization and a minimum ratio count of 2. A reverse sequence library was generated to control the false discovery rate at less than 1%.

#### 5.4. Proteomics differential abundance analysis

LFQ values were used for protein quantification. Only proteins with at least two LFQ values in at least one of the replicate groups (KO or WT) were kept for further analysis. Additionally, only proteins with more than one unique peptide detected were kept. LFQ values were log2 transformed and normalized by the median of the subgroup of proteins that were detected in the six samples. Two methods of missing value imputation were applied. When the protein was detected in two replicates, the missing value in that replicate group was defined with the k-nearest neighbors (knn) algorithm, using the average values of the five proteins (k=5) more similar (euclidean distance between LFQ values across samples) to the protein with the missing value. When the protein was detected in less than two samples in the replicate group, the missing values were defined as random values from a normal distribution with mean equal to the minimum LFQ of that sample and standard deviation equal to a global estimate made with all proteins with replicate values. It was previously checked that standard deviation estimates were not significantly affected by protein average abundance or number of detected peptides.

Statistical testing of differential abundance between the two conditions was performed with the limma R package (Ritchie et al., 2015). Adjusted p-values lower than 0.1 were considered significant (False Discovery Rate (FDR) - 10%). To avoid artifacts induced by the missing value imputation with random numbers, this imputation was repeated 30 times, and only proteins with differential abundances consistently (>50% of the times) considered statistically significant were selected for further analysis. The R code used for the analysis is available at https://github.com/GamaPintoLab/sacsin/.

#### 5.5. Functional Enrichment Analysis

We conducted Gene Ontology (GO) analysis (biological process, molecular function, and cellular component) and Kyoto Encyclopedia of Genes and Genomes (KEGG) analysis to identify biological functions and pathways enriched from the different expressed proteins. The functional enrichment analysis was conducted by using the online bioinformatics tool, the Database for Annotation, Visualization and Inte- grated Discovery (DAVID, https://david.ncifcrf.gov/). The terms with adjusted p values <0.05 were selected. The enriched functional terms were also determined by the Metascape online analytic tool (https://metascape.org/ gp/index.html#/main/step1) based on several databases, such as GO, KEGG and Reactome.

### 6. Atomic Force Microscopy

The cells were analyzed with a PicoLE Molecular Imaging system from Agilent Technologies (Keysight Technologies, Inc., Santa Rosa, CA, USA). A CP-qp-SCONT-SiO-A Nanoandmore cantilever, with nominal stiffness of 0.01 N/m and a nominal tip radius of 1 um was used in all experiments. The cantilevers are transparent to the laser except at its extremity which reduces issues with calibration. The same tip was used to measure cells of all different types to reduce bias due to different cantilever stiffness’. In total 12 cantilevers were used. To measure the mechanical properties of cells we perform grids of typically 32×32 approach/retract force-displacement curves in a range of about 30 µm. For the determination of the Young modulus of the cell we used the Hertz contact model to fit the contact portion of the approach force curve as explained elsewhere (Carapeto et al., 2020). A total of 211 cells (C6) and 168 cells (C6^Sacs-/-^) were measured.

### 7. Statistics

Statistical analysis and graphical representation of data were performed using GraphPad Prism software Version 8 (GraphPad, San Diego, CA, USA). Sample data are represented as mean ± standard error (SEM) of three independent experiments unless otherwise indicated. For statistical evaluation, one-way or two-way Analysis of Variance (ANOVA) and Tukey’s post hoc test were used for multiple comparisons. Student’s t-test was applied for comparisons in experiments with two groups. Results were considered significant when p < 0.05.

## Results and discussion

Sacsin deletion in C6 rat glioblastoma cells induces a disorganization and juxtanuclear accumulation of the three main glial intermediate filaments, vimentin, GFAP and nestin (Fig. 1)(Murtinheira et al., 2022). These alterations are also expected to extend to other central intermediate filaments and IF-associated proteins. Plectin, a crucial linker protein between IFs and various cellular components, has been identified as a sacsin interactor. Its accumulation in fibroblasts from ARSACS patients was reported, and an increased plectin signal overlapping the neurofilament bundles was observed in sacsin KO Purkinje cells from murine cerebellar sections (Bondio et al., 2023). In our C6^Sacs-/-^ model, we did not find significant changes in the expression and distribution of nuclear lamins. However, plectin showed a change in the cellular distribution in Sacs-/- cells, which was found to be absent in the juxtanuclear region where the IFs accumulate (Fig. 1A and 1D).

**Figure 1.**
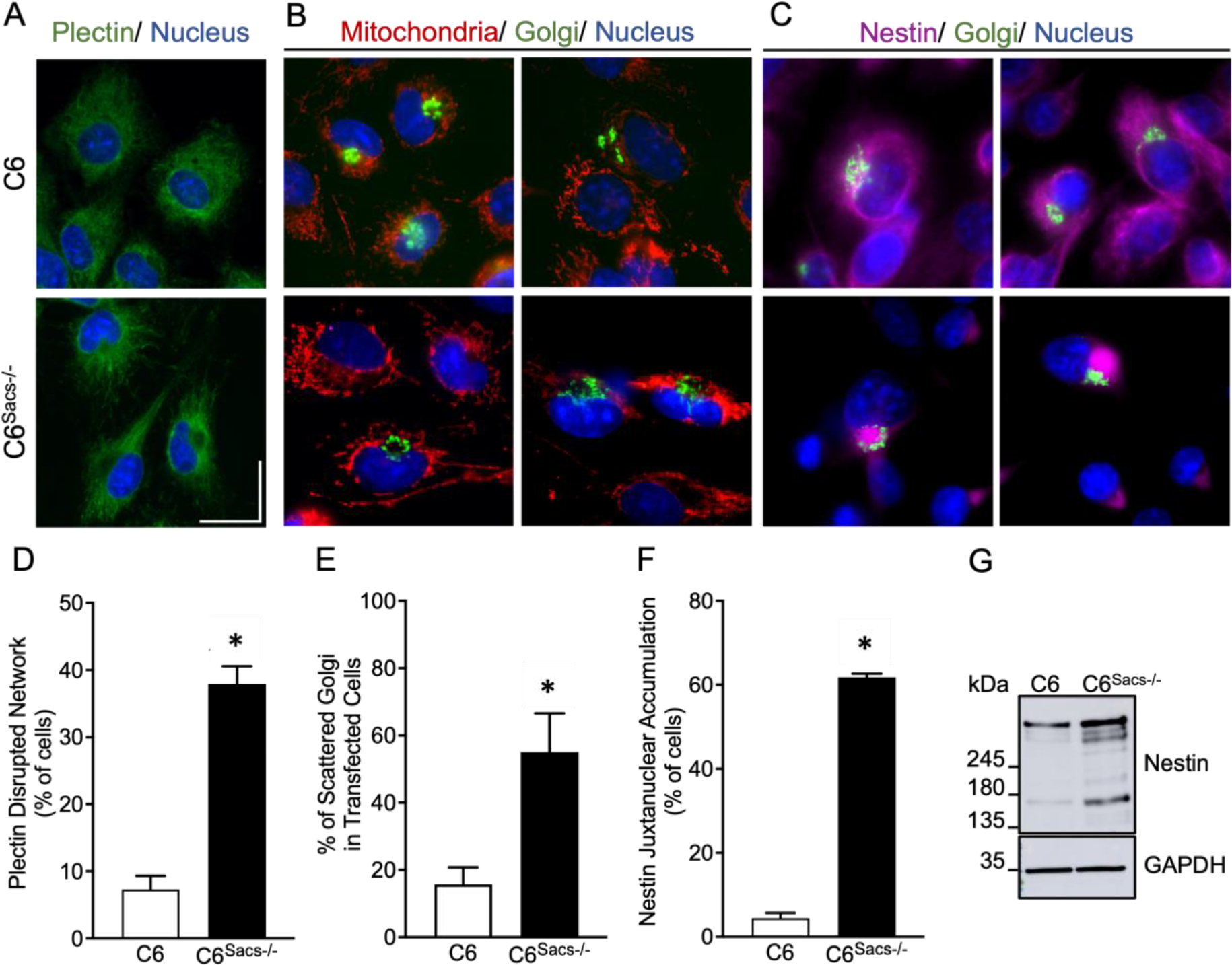
Cytoskeleton-related alterations upon sacsin deletion in C6 astroglial- like cells. Sacsin deletion disrupts the network of the cytolinker plectin in C6 cells. (A) Representative images of immunocytochemistry for endogenous plectin (green) in C6 cells and C6 ^Sacs-/-^ cells. (B-C) Representative immunofluorescence images showing the distribution of different organelles in C6 and C6^Sacs-/-^ cells transfected with EYFP-Golgi7 (green) and immunostained with an antibody against nestin (magenta). Mitochondria were counterstained with Mitotracker (red) and nuclei with Hoechst (blue). (B) Mitochondria (red) is depleted in the juxtanuclear area in C6 ^Sacs-/-^ cells, and the Golgi complex (green) is in the periphery of the same region. (C) A normal compact juxtanuclear Golgi complex (green) can be seen in C6 reference cells, whereas in C6 ^Sacs-/-^ cells, it is scattered around nestin aggregates (magenta). (D) Quantification of microscopy images from 4 independent experiments (mean ± SEM). *, significant vs. C6 reference strain (p < 0.05, Student’s t-test). In 4 independent experiments, the total number of reference C6 cells counted was 588, and the total number of C6 ^Sacs-/-^ cells counted was 995. (E) Quantification of microscopy images from 3 independent experiments (mean ± SEM). The total number of transfected cells counted was 344 in the C6 reference strain and 322 in C6 ^Sacs-/-^ cells. (F) Quantification of microscopy images from 3 independent experiments (mean ± SEM). *, significant vs. C6 reference strain (p < 0.05, Student’s t-test). In 3 independent experiments, the total number of reference C6 cells counted was 464 and the total number of C6 ^Sacs-/-^ cells counted was 697. (G) Representative Western blots showing the expression patterns of the glial intermediate filament Nestin. Scale 20 μm.

Intermediate filaments serve as scaffolds for organelles, and we and others have shown an altered mitochondrial distribution in Sacs -/- cells, moving away from the juxtanuclear IF aggregates. In our astroglial-like model, plectin exhibits a similar alteration in the cellular distribution as mitochondria (Fig. 1B). Plectin was shown to co-localize with mitochondria in striated muscle (Reipert et al., 1999) or directly bind with both vimentin and mitochondria (Winter et al., 2008). In neurons, plectin dysfunction was shown to alter microtubules dynamics and increase tau accumulation (Valencia et al., 2021).

Previous reports also indicated altered Golgi distribution in non-glial models of ARSACS, and we confirmed this alteration in C6^Sacs-/-^ cells. In C6 cells, the Golgi apparatus exhibits a compacted juxtanuclear localization. Upon sacsin deletion, the Golgi apparatus becomes dispersed, distributing around IF aggregates (Fig 1, B, C, D), similar to what happens in ARSACS patientś fibroblasts (Duncan et al., 2017).

Intermediate filaments are essential for the maintenance of mechanic and viscoelastic properties of cells, and Atomic Force Microscopy (AFM) confirmed a decrease in the average height of the perinuclear region and a small but consistent decrease in the apparent Young modulus, indicating structural alterations in cell bodies (Fig. 2). We assume the elasticity modulus is log normally distributed. A comparison between the two distributions results in statistically different elasticity modulus of 163 Pa and 115 Pa respectively for the C6 and C6^Sacs-/-^. The reduction of this parameter in sacsin knockout cells can be associated with alterations in internal structure and cytoskeleton organization, confirming a loss of cell integrity and higher susceptibility to deformation.

**Figure 2.**
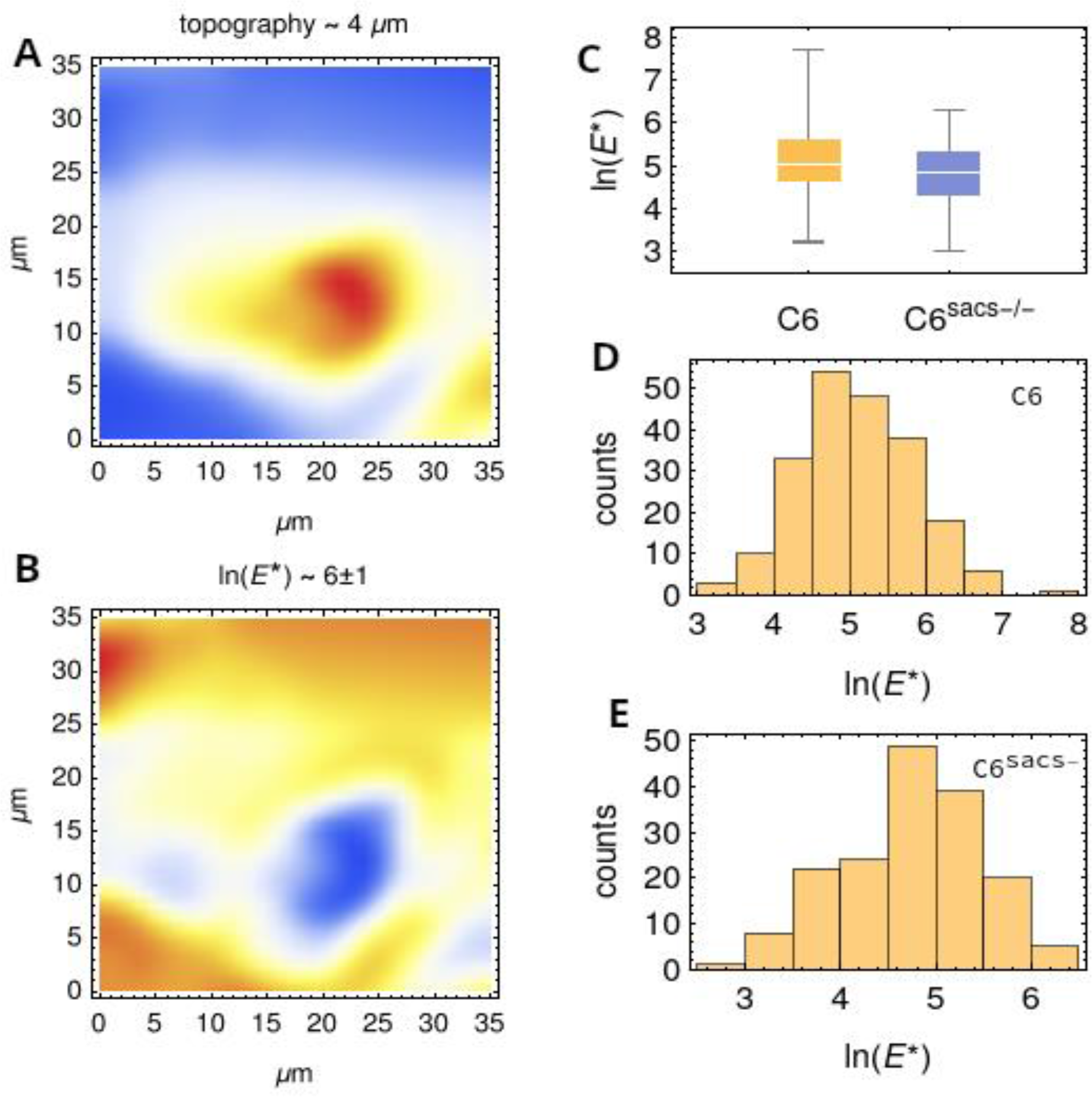
Mechanical alterations caused by sacsin deletion and intermediate filament disorganization in C6 astroglial-like cells. (A-B) representative plots of the topography and elasticity maps obtained from a 32⨉32 grid (the color range is indicated on top). (C) Box whiskers chart of C6 and C6 **^Sacs-/-^**. (D-E) Histograms of the logarithm of the apparent Young modulus for C6 and C6Sacs-/- cell respectively.

To understand global molecular and cellular changes in C6 cells by loss of sacsin, we carried out a proteomic analysis on total protein extracts from C6 and C6^Sacs-/-^ cells. For each cell line, we conducted three biological replicates. We then selected proteins that were identified in at least two replicates. This approach identified 1552 proteins; the VENN diagram showed 1490 common to both datasets, 103 unique to the C6 reference cells and 30 exclusive to Sacs-/- cells (Fig. 3, A). Differentially expressed proteins are shown as a volcano plot where blue dots and red dots represent sets of proteins expressed at significantly lower and higher levels in Sacs-/- cells, respectively (Fig. 3, B). The selection criteria for differentially expressed proteins (DEPs) were set at a two-fold change and statistical significance (p < 0.05) between the Sacs-/- model and reference cells. The expression of 140 proteins significantly differed between reference and C6^Sacs-/-^ cells; 104 were upregulated, and 36 were downregulated (Fig. 3, B). Through DAVID analysis, the gene ontology (GO) analysis results showed that biological process terms were enriched in terms mainly related to adhesion, trafficking and migration, as well as a cellular response to mechanical stimulus, intercellular signalling pathways and cell redox homeostasis (Fig. 3, C). According to the cellular component GO enrichment, the DEPs play a role in structural and cytoplasmic integrity (cytoplasm, cell membrane, cytoskeleton, nucleoplasm and stress fibres) and cellular connection and communications (synapse, cell-cell adherent junction, focal adhesion proteins and podosome) (Fig. 3, D). These results are consistent with disorganization of the IF networks and alterations in the mechanical and viscoelastic properties of cells observed by AFM. Actin binding and protein tyrosine kinase binding were two molecular function-enriched terms for the DEPs (Fig. 3, E). The pathways associated with the DEPs were identified using the Metascape platform and the Molecular Complex Detection (MCODE) algorithm was then applied to identify neighborhoods where proteins are densely connected. Four independent clusters were identified (Fig. 3, F). The significantly enriched functional descriptions for each complex are as follows: MCODE1 include rRNA processing (0006364), rRNA metabolic process (GO:0016072), ribosome biogenesis (GO:0042254); MCODE2 include positive regulation of cell-substrate adhesion (GO:0010811), focal adhesion (rno04510), Rap1 signaling pathway (rno04015); MCODE3 include cell junction assembly (GO:0034329), cell junction organization (GO:0034330). Gene ontology and functional terms analysis confirmed substantial alterations in cellular and molecular functions related to the cytoskeleton. Additionally, we found dysregulation of signaling pathways supporting the idea that intermediate filaments are scaffolds relevant to cell signaling.

**Figure 3.**
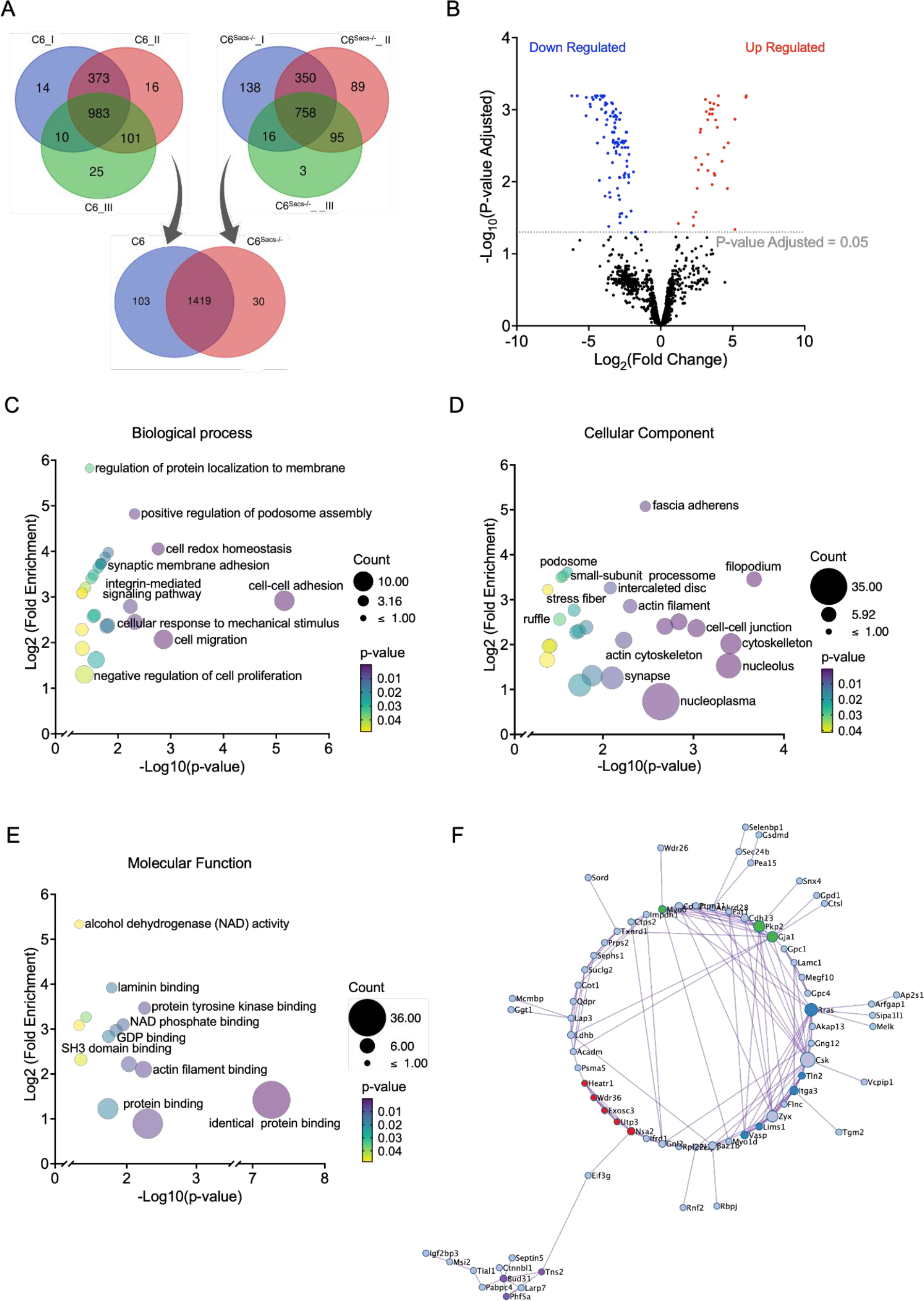
Sacsin deletion induces alterations in cell functions associated with cell motility and mechanics. (A) Volcano plot showing protein abundance differences between C6 **^Sacs-/-^** cells and C6 reference strain. Black points represent unchanged proteins, and red and blue represent the upregulated and downregulated proteins in C6**^Sacs-/-^** cells, respectively; the horizontal dashed line indicates adjusted p-value = 0.05. (B) Venn diagram illustrating the number of proteins expressed by three biological replicates of C6 reference strain and C6 **^Sacs-/-^** cells. The overlapping areas between the biological replicates indicate the numbers of commonly expressed proteins for each cell type. Venn diagram showing the number of proteins expressed by each cell line. The overlapping areas indicate the number of proteins commonly expressed by the C6 reference strain and C6 **^Sacs-/-^** cells. (C-E) Gene Ontology analysis of differentially expressed proteins in C6 **^Sacs-/-^** vs C6 using DAVID software. Detailed information relating to changes in the biological process (C), cellular component (D) and molecular function (E). (F) Protein-protein interaction enrichment analysis. The network includes proteins interacting physically with at least one other member from the input list. MCODE algorithm identifies nodule density and assigns a unique colour to each network. Relevant statistical analysis is presented as a log10(p) value. DAVID: Database for Annotation, Visualization and Integrated discovery; GO: Gene Ontology. MCODE: Molecular Complex Detection.

Proteomics analyses of C6 and C6^Sacs-/-^ protein extracts also suggested possible alterations in the levels of Endoplasmic reticulum oxidase 1 (ERO1), NAD(P)H:quinone oxidoreductase (NQO1), Glutaredoxin 3 (GLRX3) and E3 ubiquitin ligase (Ube3A) which we tried to confirm individually by immunoblotting (Fig. 4). We confirmed reduced levels of ERO1 and NQO1 upon sacsin deletion in C6 cells but did not find significant alterations in GLRX3 levels. ERO1, a thiol oxidase, is essential for the oxidative process of protein folding within the ER, and an increase in oxidative protein folding can lead to generation of ROS in the ER (Bhandary et al., 2012). ERO1 is overexpressed in cancer and altered in diabetes and neurodegenerative diseases (Shergalis et al., 2020). The reduction of ERO1 levels observed in C6^Sacs-/-^ cells can also disrupt the redox equilibrium and accumulate misfolded proteins in the ER. Proteomic analysis in other ARSACS models showed involvement of ER calcium-binding proteins associated with protein folding quality control, such as the chaperone calreticulin (CALR) (Morani et al., 2022). Additionally, impaired ER and mitochondrial trafficking to distal dendrites may be the cause of Ca^2+^ dysregulation in these locations(Bondio et al., 2023). These observations suggest a connection between the ER and ARSACS. NQO1 is a chemoprotective enzyme against the harmful oxidative effects of quinones (Ross & Siegel, 2021). The absence of this enzyme can result in alterations in intracellular redox state and seizures, suggesting an important role of the protein in the CNS. Neurons do not contain significant amounts of this protein, but it is relatively abundant in glial cells (Stringer et al., 2004). NQO1 was also reduced in another ataxia, SCA17, but increased in patients with Alzheimer’s disease (Lee et al., 2014; SantaCruz et al., 2004).

**Figure 4.**
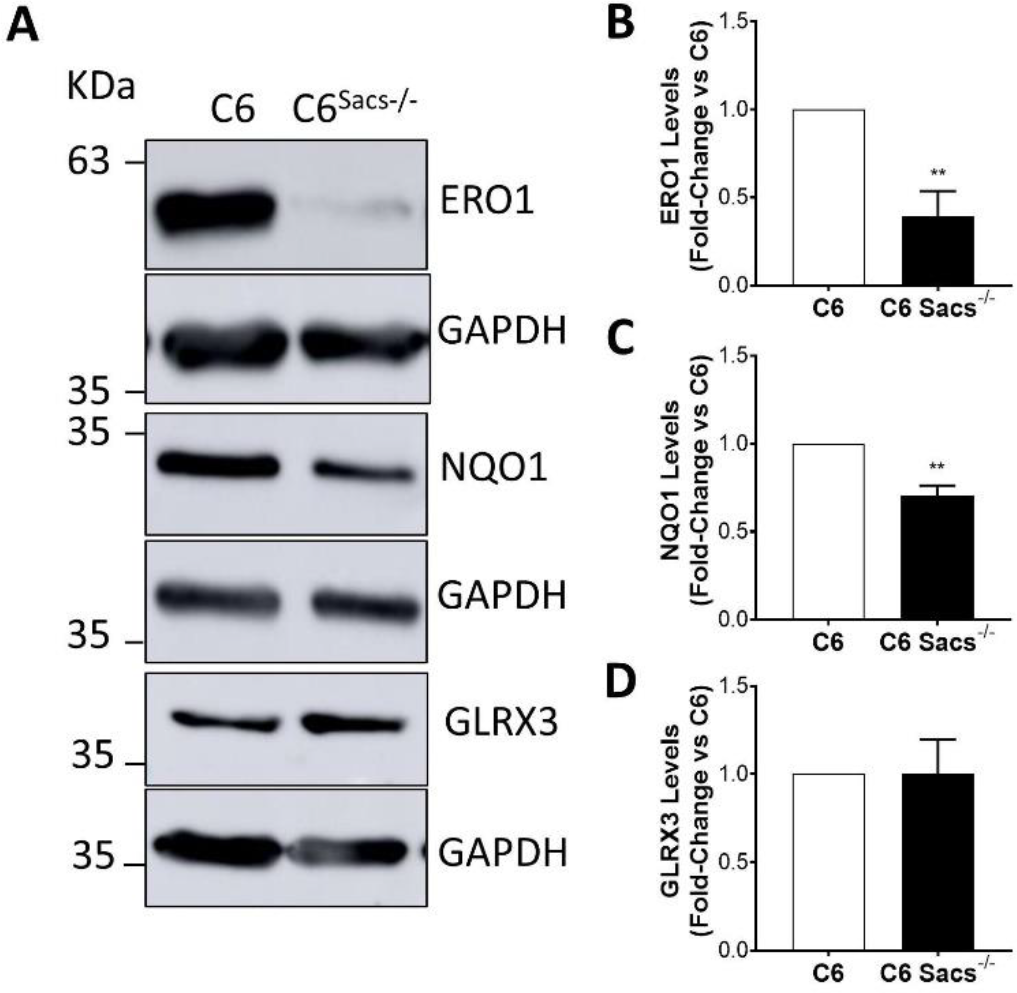
Sacsin deletion induces alterations in the levels of key regulators of stress-associated pathways. Representative western blots showing ERO1, NQO1 and GLRX3 expression levels and densitometric analysis of western blots from at least 3 independent experiments normalized versus GAPDH. (A) Representative western blots. (B-D) Densitometric analysis of bands, normalized versus GAPDH. Data are shown as mean ± SEM; unpaired student’s t-test: **P < 0.01.

We have previously shown that C6^Sacs-/-^ cells show alterations in cytokine pathways relevant to development and neuroinflammation, such as STAT3 (Murtinheira et al., 2022). Smad1 is a transcription factor that mediates TGF-beta and Bone Morphogenetic Proteins (BMPs), and that is relevant for development and neuroinflammation. We and others have shown that Smad1 also forms complexes with STAT3, bridged by p300/CBP, and both pathways can have synergistic effects in the expression of glial-specific genes, such as GFAP (Herrera et al., 2010). We observed that the levels of Smad1 were significantly decreased in C6^Sacs-/-^ cells (Fig. 5).

**Figure 5.**
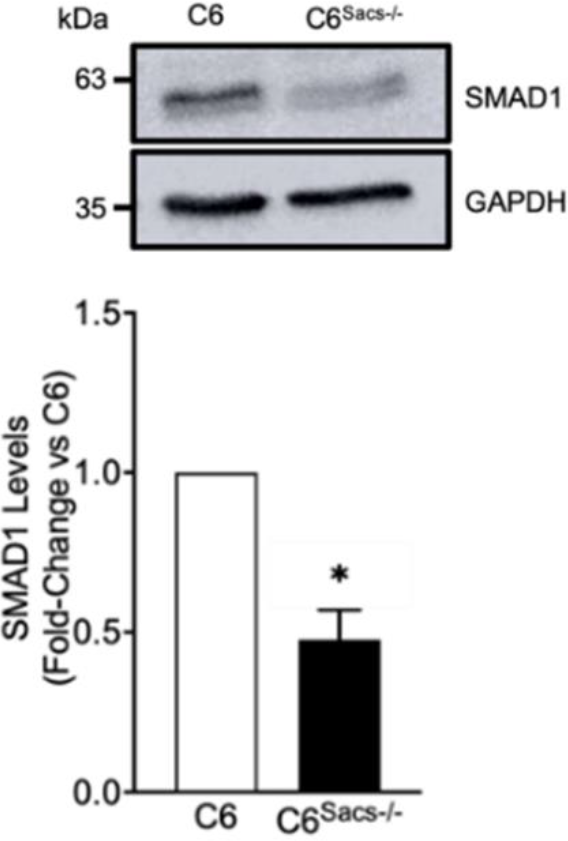
Sacsin deletion induces a decrease in the levels of Smad1, a transcription factor relevant for development and neuroinflammation. Representative western blots showing Smad1 expression levels and densitometric analysis of western blots from at least 3 independent experiments normalized versus GAPDH. Data are shown as mean ± SEM; unpaired student’s t-test: *P < 0.05.

Cellular and animal models of disorders associated with alterations in glial IF networks, such as Alexander disease and Giant Axonal Neuropathy, often show an increased sensitivity of cells to stress and injury. Consistently, C6^Sacs-/-^ cells displayed higher sensitivity to rotenone-induced oxidative stress (Murtinheira et al., 2022), and we observed that this sensitivity could be extended to other forms of stress, such as serum deprivation or exposure to radiation (Fig. 6). Serum deprivation does not substantially change the morphology of reference C6 cells but leads to a round morphology in C6^Sacs-/-^ cells (Fig 5, A). Serum deprivation leads to a reduction in cell number in both C6 and C6^Sacs-/-^ cells but the reduction is significantly higher in Sacs-/- cells (Fig. 6, B). The generation of reactive oxygen species (ROS) upon serum starvation was significantly increased in Sacs-/- cells compared to reference cells (Fig 6, C).

**Figure 6.**
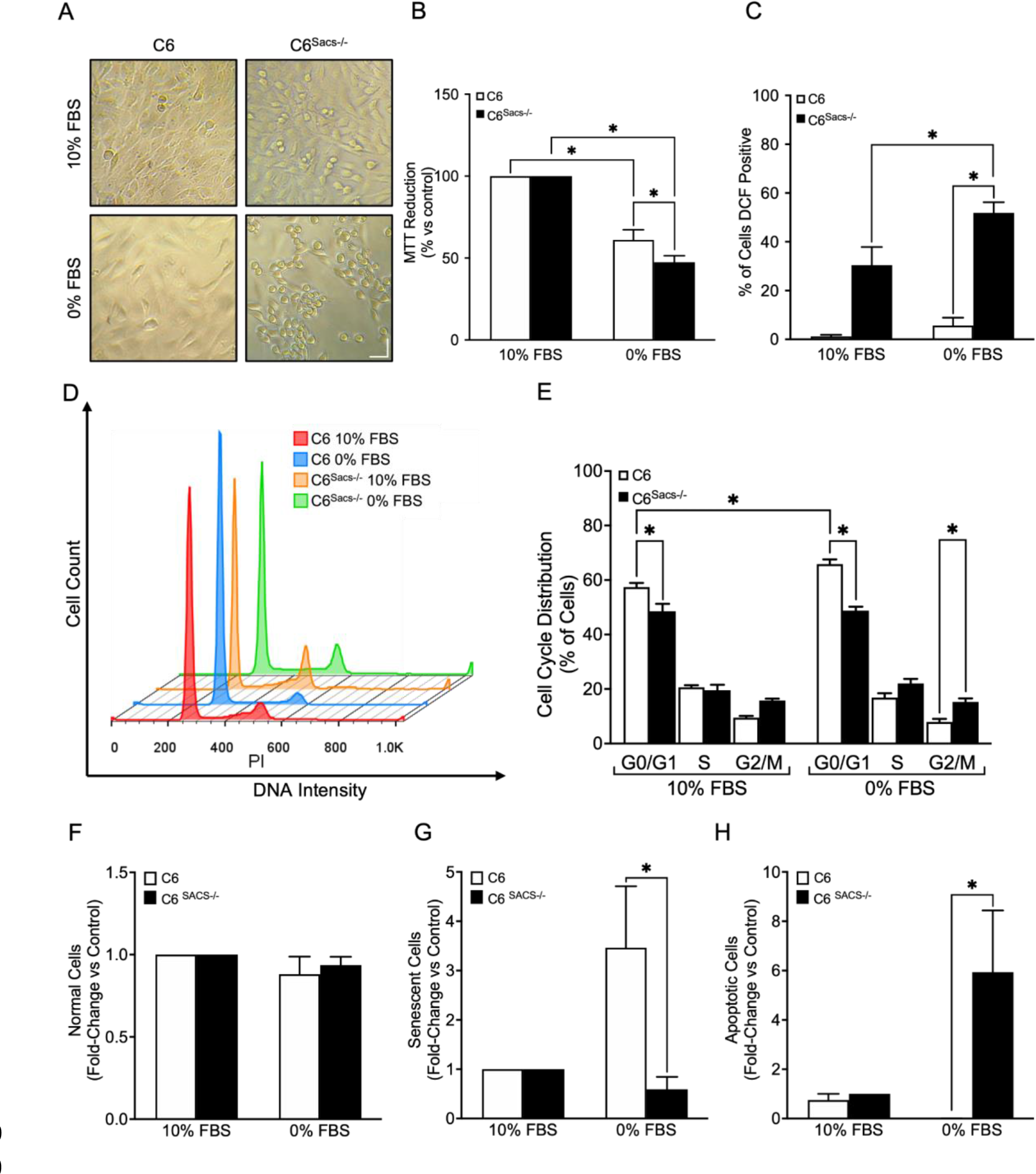
Sacsin-knockout cells have higher sensitivity to serum starvation. Serum starvation reduces cell viability in C6 **^Sacs-/-^** cells. C6 and C6 **^Sacs-/-^** cells were grown with or without 10% FBS for 48h. (A) Representative bright-field images of cell morphology and density under the serum starvation stress. (B) Cell viability was analysed using the MTT assay. A significant decrease in the percentage of viable cells was observed in C6 and C6 **^Sacs-/-^** compared to controls. (C) Intracellular ROS levels were monitored by DCFDA staining followed by flow cytometry. Plot represents percentage of DCF-positive cells. In serum-starved C6 **^Sacs-/-^** cells, the percentage of DCF-positive cells increased significatively compared to control (D-E). (D) Cell cycle distribution was analysed by flow cytometry. Representative histogram of the gated cells in the G0/G1, S, and G2/M phases. (E) Quantitative analysis of distribution of the cells in each phase was performed from at least 10,000 cells per sample. (F-G) Hoechst-stained cell nuclei were subjected to nuclear morphometric analysis using the ImageJ NII_Plugin. The bar graphs indicate relative fold changes for senescent (F) and apoptotic cells (G) compared to control cells. Data are represented as the mean ± SEM (standard error in the mean). Statistical significance was determined by One-way analysis of variance (ANOVA), followed by Holm-Šídák’s multiple comparisons test. The differences were considered statistically significant when p < 0.05.

Stress conditions can impact the cell cycle dynamics, as C6 cells show an accumulation of cells in the G0/G1 phase upon starvation (Fig. 6, D, E). However, C6^Sacs-/-^ cells exhibit a decrease in the proportion of cells in G0/G1 phase and a higher proportion of cells in G2/M phase compared to reference cells, regardless of serum availability (Fig. 6, E). These findings collectively suggest an induced cell cycle arrest in response to stress stimuli. Morphological analysis of nuclei (Filippi-Chiela et al., 2012) indicated that serum starvation increased the number of senescent C6 cells (Fig 6, F), while inducing a significant rise in the number of apoptotic C6^Sacs-/-^ cells (Fig 6, G).

HMC3 is a microglial cell line that was isolated from human brain that also expressed sacsin, just as N9 rodent microglial cells (Murtinheira et al., 2022). We aimed to develop a human microglial model of ARSACS, deleting sacsin in HMC3 cells using the CRISPR/Cas9 approach previously described (Murtinheira et al., 2022). After selecting a clone that did not express detectable levels of sacsin, we evaluated whether this model was similar to the C6 model of ARSACS. As already described, sacsin deletion induces the accumulation of intermediate filaments in the juxtanuclear region, with the concomitant depletion of mitochondria in the same region. In the C6 astroglial-like model, this subcellular redistribution is accompanied by higher overall levels of IFs. Although we did not see higher levels of vimentin in HMC3^Sacs-/-^, juxtanuclear accumulation of the protein is similar to what we have previously described in C6 cells (Fig 7A-C).

**Figure 7.**
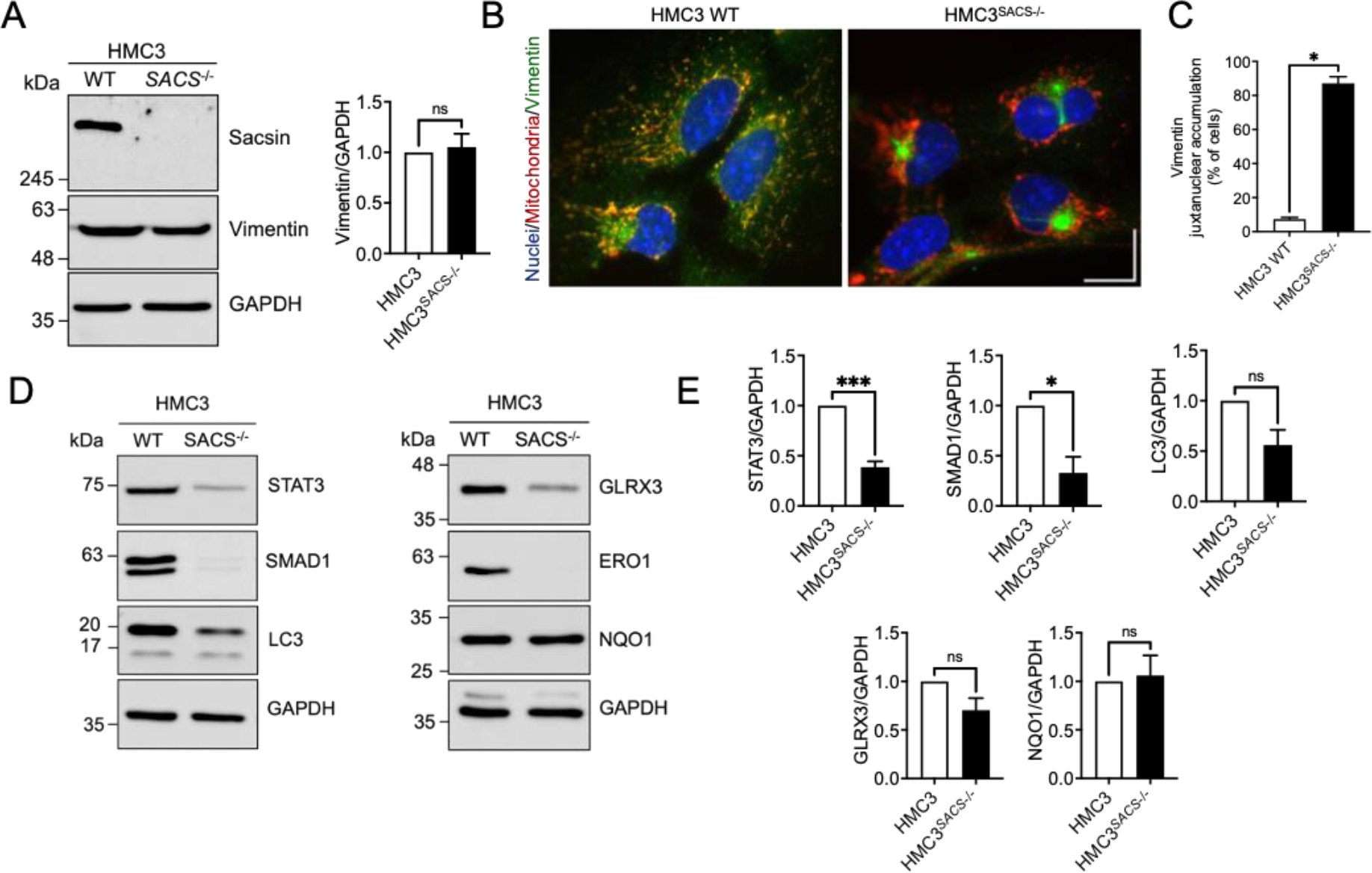
Sacsin deletion in human microglial cells disrupts vimentin networks and developmental and inflammatory-related pathways. (A) Representative Western blots of total cell lysates from HMC3 human microglial cells and an HMC3 **^Sacs-/-^** clone showing the expression patterns of sacsin and vimentin. Vimentin bands from 3 independent experiments (n=3) were quantified by densitometry, and results were normalized for GAPDH. Results represented as mean±SEM. (B) Representative immunocytochemistry images showing the distribution of vimentin (green); mitochondria (Mitoview, red) and nuclei (Hoechst, blue) in HMC3 and HMC3 **^Sacs-/-^** cells. Scale bar 20 µm. (C) Quantification of microscopy images from 3 independent experiments (mean±SEM). *, significant vs. HMC3 reference strain (P value=0.0226, Welch’s t-test). The total numbers of HMC3 reference cells and HMC3 **^Sacs-/-^** counted in the 2 independent experiments were 777 and 831, respectively. (D) Representative Western blots showing the expression patterns of SMAD1, STAT3, LC3, GLRX3, ERO1 and NQO1. (E) Densitometric analysis of western blots from at least 3 independent experiments normalized for GAPDH. Results represented as mean±SEM. *, significant vs. HMC3 reference strain (SMAD1: P value=0.0257; STAT3: P value=0.0004 Welch’s t-test); ns, non-significant.

In our previous study, we demonstrated that sacsin deletion in C6 cells leads to alterations in the response to inflammatory cytokines, decreasing the levels of STAT3 protein. In the HMC3 model we also observed a significant reduction in ERO1, STAT3 and Smad1 levels upon sacsin deletion (Fig 7D-E). Autophagy-associated protein LC3 and GLRX3 show a non-significant tendency towards reduction, while NQO1, which decreased upon sacsin deletion in the C6 model, did not show alterations in the HMC3 model. In summary, there are Next, we assessed the function of HMC3^Sacs-/-^ cells by addressing their phagocytosis ability (Fig. 8). We used two types of phagocytosis inducers: myelin debris to assess microglia neuroprotective ability to clear the CNS debris, and zymogen-coated beads to assess microglia innate immune response. We were able to show that HMC3^Sacs-/-^ cells maintain their phagocytic ability, but with a higher variability than the HMC3 cells. Indeed, HMC3^Sacs-/-^ cells phagocyted a significantly higher amount of myelin debris and a slightly higher number of zymogen beads than the HMC3 ones (Fig. 8A-B), suggesting a potentially increased ability for clearance and pathogen reactivity. Reactive microglia usually shift their morphology into a more spherical volume, so we next evaluated cell sphericity and surface area to volume ratio (Fig. 8C-D). Interestingly, while HMC3 cells react to both myelin debris and zymogen beads altering their shape into a more spheroid morphology and showing a lower surface area to volume ratio (in accordance with a more reactive morphology), HMC3^Sacs-/-^ cells fail to change their morphology upon stimulation, suggesting that an altered cytoskeleton response may be present.

**Figure 8.**
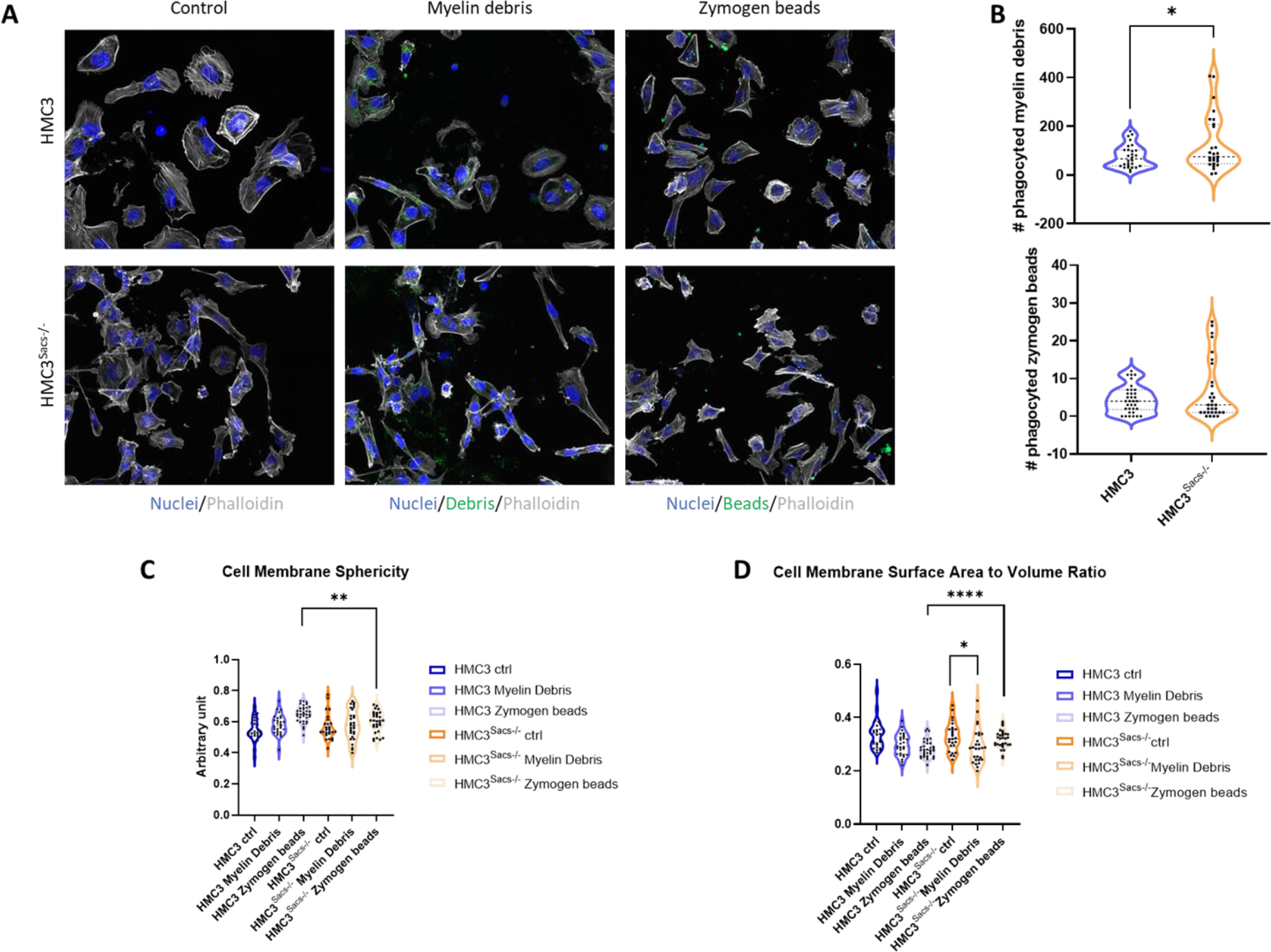
Alterations in the phagocytosis functions of HMC3^Sacs-/-^ cells. (A) representative images of HMC3 human microglial cells and an HMC3 **^Sacs-/-^** clone showing their ability to phagocyte both myelin debris (green) or zymogen beads (green). (B) Quantification of myelin debris or zymogen beads phagocyted by HMC3 and HMC3 **^Sacs-/-^** clone from 3 independent experiments. *, significant vs. HMC3 reference strain (*P value<0.05). The total numbers of HMC3 reference cells and HMC3**^Sacs-/-^** counted in the 3 independent experiments were 40. Quantification of (C) cell membrane sphericity and (D) surface area to volume ratio of HMC3 reference cells and HMC3 **^Sacs-/-^** using AIVIA software (**P value<0.01, ***P value<0.001, ****P value<0.0001). The total numbers of HMC3 reference cells and HMC3 **^Sacs-/-^** counted in the 3 independent experiments were 40.

## Conclusions

Our results support the idea that alterations in the IF and mitochondrial networks in glial cell models of ARSACS have biologically relevant consequences on the mechanical and viscoelastic properties of cells, their susceptibility to stress and their inflammation-related signalling pathways. These 3 properties of glial cells are key for a coordinated response to CNS insults during neuroinflammation, and point at a possible role for glial cell dysfunction in ARSACS. Further analysis on glial-specific functions of sacsin should be done to determine the extent of their contribution to ARSACS symptoms and histopathological features.

## Acknowledgements

We acknowledge support from the BioISI/FCUL Microscopy Facility, a node of the Portuguese Platform of BioImaging (PPBI-POCI-01-0145-FEDER-022122) and the BioISI Mass Spectrometry Facility, with support from the Portuguese Mass Spectrometry Network, integrated in the National Roadmap of Research Infrastructures of Strategic Relevance (ROTEIRO/0028/2013; LISBOA-01-0145-FEDER-022125). We would also like to thank the Flow Cytometry Facilities at Instituto Gulbenkian de Ciência (Oeiras, Portugal) and Instituto de Medicina Molecular João Lobo Antunes (Lisbon, Portugal) for their technical support. FH, VMT, FRP and MSR were supported by centre grants https://doi.org/10.54499/UIDP/04046/2020 and https://doi.org/10.54499/UIDB/04046/2020 (BioISI Research Unit) and individual grants (Refs. PTDC/BTM-TEC/28554/2017 and https://doi.org/10.54499/PTDC/FIS-MAC/2741/2021 to VMT and MSR, respectively) through Fundação para a Ciência e Tecnologia. FH and AF were supported by a grant from the ARSACS Foundation (Canada), which included two fellowships to ASB and LM. FM was supported by FCT (Ref. SFRH/BD/133220/2017). AF, EF were supported by centre grants UIDB/04138/2020 and UIDP/04138/2020 from FCT to iMed.ULisboa. Funded by the European Union (TWIN2PIPSA, GA 101079147). Views and opinions expressed are however those of the author(s) only and do not necessarily reflect those of the European Union or European Research Executive Agency (REA). Neither the European Union nor the granting authority can be held responsible for them.

## Contributions

The HMC3 cell line was developed by FM, LM and ASB, who also carried out immunocytochemistry, protein extraction and immunoblotting experiments in both C6 and HMC3 cells, with punctual contribution by PN. JB, TTR and MSR performed AFM experiments. FM prepared the protein samples for proteomics, VMT was responsible for mass spectrometry analysis, and FRP supervised the computational analysis of proteomics data, mainly carried out by FM. EF and AF characterized the morphology and phagocytic activity of HMC3 cells. FM, ASB, AF, VMT, FRP, MSR and FH contributed to the preparation of figures, writing and revision of the manuscript. MSR and FH designed the experiments, coordinated the teams, and provided the funding for most research activities.

